# Aberrant inheritance of extrachromosomal DNA amplifications promotes cancer evolution

**DOI:** 10.1101/2025.09.19.677276

**Authors:** Shir Marom, Inbar Lifshits Dayan, Venkata Narasimha Kadali, Mădălina Giurgiu-Kraljič, Gabriela Koifman, Karen Hakeny, Madhuri Chaurasia, Orléna Benamozig, Reinat Nevo, Ido Azuri, Julia Ryvkin, Ron Rotkopf, Gil Stelzer, Nino Oniashvili, Jacques Mardoukh, Sarah Pollock, Nika Iremadze, Zohar Shipony, Meital Kupervaser, Noa Wigoda, Dena Leshkowitz, Zvulun Elazar, Esther Berko, Anton G. Henssen, Ofer Shoshani

## Abstract

Gene amplification in the form of extrachromosomal DNA (ecDNA) is a frequent driver in multiple cancer types. As ecDNA lack centromeres, their mitotic segregation does not follow traditional inheritance principles. However, the mechanisms that govern ecDNA fate following mitosis remain unclear. We found that ecDNA undergo numerical and structural optimization under increased selective pressure, with mitotic chromosomal tethering, or detachment, dictating ecDNA fate. When tethered, ecDNA aggregates promote uneven distribution into the newly formed daughter cells, thereby driving inter-cellular numerical heterogeneity and rapid increase of amplification under selective pressure. Mitotically detached ecDNA frequently encapsulate within micronuclei of variable size and content that appear to be highly fragile. Strikingly, ecDNA enclosed in very small micronuclei, which we term nanonuclei, are being actively degraded through autophagy. Together with ongoing structural rearrangements, nanonuclear ecDNA degradation promotes their structural evolution, which facilitates cancer cell adaptation. Our work highlights ecDNA aggregation, micronucleation, and degradation, as pivotal events in directing cancer genome evolution trajectories.

## Main

Gene amplification is a major driver in multiple cancers and is associated with poor prognosis^1–7^. The two distinct types of amplification, intrachromosomal (homogenous staining regions, HSRs) and extrachromosomal (extrachromosomal DNA, ecDNA, also known as double minute chromosomes) differ both in their mechanism of formation and their maintenance within cancer cells^5,8–13^. The acentric nature of ecDNA distinguishes it from canonical chromosomal inheritance, leading to random mitotic distribution and subsequent increase in intra-tumor heterogeneity^14–16^. The capacity of ecDNA to tether to chromosomes^17–19^ is often disrupted, leading to their detachment and missegregation into micronuclei^11,19,20^. As chromosome micronucleation often leads to chromothripsis^21–23^, the shattering and random re-ligation of chromosomes^12^, it was proposed that ecDNA might be subjected to successive chromothriptic events as cancer cells propagate^11,24^. Indeed, ecDNA frequently present complex numerical and structural alterations, also during adaptation to therapeutic interventions^2,11,14,25–33^. Finally, ecDNA clustering through the binding of the bromodomain and extraterminal domain (BET) protein BRD4 was suggested to promote transcriptional output^34^ and impact the co-inheritance of multiple ecDNA species^2^, thus affecting the evolutionary trajectories of ecDNA under therapy. Here, we follow the fate of ecDNA during and following mitosis to reveal the mechanisms underlying ecDNA evolution that promote cancer cell adaptation.

## ecDNA optimization drives cancer adaptation

To model the evolution of ecDNA under increased selective pressure, we created an isogenic cohort of HeLa clones with increasing levels of methotrexate (MTX) resistance driven by *DHFR*+ ecDNA amplification. We subjected three independent clones^11^-that harbor *DHFR*+ ecDNA of variable sizes (clone 1 – 2.1 Mb, clone 2 – 7.43 Mb, and clone 3 – 2.19 Mb) and that are initially resistant to 40nM MTX, to step-wise increase of MTX concentrations up to 10,000 nM (250-fold change). We examined 1,057 mitotic spreads using DNA-FISH (fluorescence in situ hybridization, Fig. 1a) to measure *DHFR*+ ecDNA amounts at each MTX concentration (Fig. 1b, total of 118,128 ecDNAs scored). Exposure to elevated MTX concentrations led to an average increase of ecDNA content by over 36-fold, as also determined using deep (58X) short-read whole genome sequencing (WGS) (Extended Data Fig. 1a). EcDNA copies were highly variable at each MTX dose: while cells resistant to 10,000 nM of MTX had an average of 362±323 ecDNA per cell, some cells contained over 2,000 ecDNA and others had less than 100 (Fig. 1b). This large heterogeneity is seen in the ecDNA distributions of all three clones (Extended Data Fig. 2). Of relevance, although the average content of ecDNA increased from 10±13 to 74±80 during the transition from 40 to 1,600 nM MTX resistance, a subset of cells within this concentration window lacked ecDNA altogether. Examination of the *MYCN+* ecDNA neuroblastoma cell-line CHP-212, revealed it harbors an average of 180±117 ecDNA per chromosome spread (Extended Data Fig. 1b-c), with one spread showing over 500 ecDNA and others having less than 50, reflecting a 10-fold difference within this cell line. Such high and variable ecDNA content was also observed in Colo320DM, a colorectal cancer cell-line, harboring *MYC+* ecDNA (180±128 ecDNA/spread), and UACC-1598, an ovarian cancer cell line with *MYCN*+ ecDNA (95±62 ecDNA/spread), but not in the small cell lung cancer cell line NCI-H524 in which 24±23 large *MYC*+ ecDNA were found on average (Extended Data Fig. 1b,d). Using the amplicon repository database ^35^, we found that NCI-H524 has ∼200 copies of *MYC*, indicating that each ecDNA in this cell line contains 8-9 copies of the *MYC* gene. Of relevance, SNU-16, a gastric cancer cell-line, harboring both *MYC* and *FGFR2* amplification in ecDNA was previously reported to show great heterogeneity as well: 5-300 copies of *MYC* and 100-500 copies of *FGFR2*^2^. To examine ecDNA content and heterogeneity in clinical settings, we performed DNA-FISH on tumor tissue, bone marrow aspirations, and cerebral spinal fluid (CSF), obtained from neuroblastoma patients positive or negative for *MYCN* ecDNA. Strikingly, ecDNA+ patient samples exhibited high amplification levels (49±51 ecDNA per cell), showing great variability, with the highest ecDNA cellular count of 458 copies, 9-fold increase above the average (Fig. 1c-d, Extended Data Fig. 1e-f), similarly to the examined cancer cell lines. Importantly, as tumor sections often contain non-cancer cells that lack ecDNA, it convolutes ecDNA scoring, thus leading to significant underestimation.

**Figure 1:**
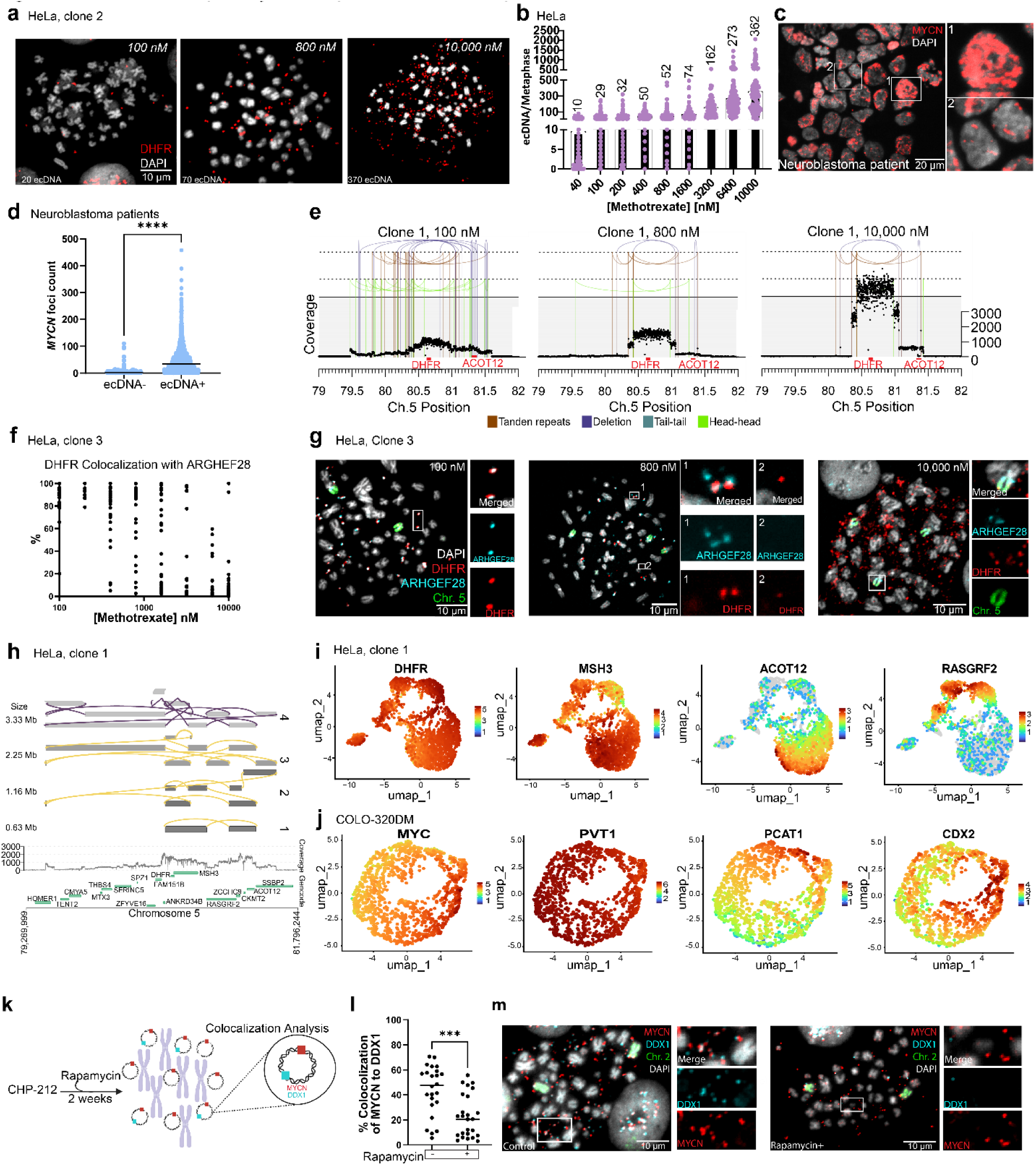
Numerical and structural plasticity of ecDNA promotes cancer cell adaptation. a, Representative DNA-FISH images of metaphase spreads from ecDNA+ HeLa clone 2. Cells shown with average amount of ecDNA amplification in methotrexate (MTX) resistant clones initially harboring DHFR+ ecDNA in three different resistance concentrations: 100 nM, 800 nM and 10,000 nM. b, Quantification of ecDNA amplification in MTX resistant clones initially harboring *DHFR+* ecDNA (n=3 clones, N=1,057 metaphase spreads, total of 118,128 ecDNA were counted) challenged with increasing doses of MTX. Amplification content was determined using DNA-FISH as shown in panel a. c, Representative DNA-FISH image of neuroblastoma patient tumor section exhibiting high amplification and heterogeneity of *MYCN+* ecDNA. d, Quantification of *MYCN* foci in neuroblastoma patients (n=3 ecDNA*-*, 3 ecDNA*+*). ecDNA*+* patients showed an average of 55±51 ecDNA, ecDNA-patients showed 3±7. Welch’s t-test two-tailed. e, Copy number and structural variation profiles of HeLa clone 1 at three different MTX resistance concentrations (100/800/10,000nM). f, Percent of colocalization of *DHFR* and *ARHGEF28* genes per cell in HeLa clone 3 at increasing resistance of MTX. X axis is logarithmic scale. g, Representative DNA-FISH images of HeLa clone 3 resistant to increasing MTX resistance concentrations showing colocalization, or lack thereof, of the *DHFR* and *ARHGEF28* genes. h, Reconstruction of ecDNA structures from HeLa clone 1 using structural variant consensus calls from long- and short-read data analyzed with Decoil^36^. The top four amplicon structures are presented (structure proportions are 1-305, 2-160, 3-92, 4-44, and are indicated by the shades of grey colors). i, UMAP plots from single-cell RNA sequencing analysis of HeLa clone 1. The expression of four different genes found on the amplicon are presented (from left to right): *DHFR*, *MSH3*, *ACOT12* and *RASGRF2*. Expression levels are presented using the indicated color scales (red – high, blue – low expression). j, UMAP plot single-cell RNA-seq of Colo320DM showing four different variable genes that are found on the amplicon (from left to right): *MYC*, *PVT1*, *PCAT1* and *CDX1*. Each color describes how highly expressed the gene is within the cell. k-m, (k) Illustration of experimental set-up. CHP-212 cells were treated with rapamycin for 2 weeks (DMSO was used as a control) and metaphase spreads were prepared for DNA-FISH. (l) Quantification of *MYCN* colocalization with *DDX1* (Welch’s t-test one-tailed). One of three experiments is presented (representative DNA-FISH images presented in panel m, see Extended Data Figure 1q-r for experiments 2 and 3).

Our analyses reveal that oncogene ecDNA amplification in cell lines or patient samples exhibits comparable ecDNA amount and heterogeneity to HeLa cells resistant to high (3,200-6,400 nM) MTX concentration, thus representing highly evolved ecDNA. Previous comparison of pre-cancer and esophageal adenocarcinoma samples revealed significant increase in ecDNA structural heterogeneity suggesting tumor cells contain evolved ecDNA species^1^. Indeed, a pan-cancer analysis found that ecDNA are characteristic of advanced tumors and their structural complexity and size are preserved over time, suggesting that already upon initial diagnosis, ecDNA present a highly evolved structure^7^. To study earlier ecDNA structural evolution, we used WGS to sequence samples from two of the HeLa clones at each MTX concentration step (Fig. 1e and Extended Data Fig. 3). Under increasing MTX pressure, while both clones had significant increase in *DHFR* reads, the amplicon size in clone 1 decreased from ∼2Mb to ∼0.5Mb and showed reduced complexity (Fig. 1e and Extended Data Fig. 1g). Although amplicon complexity was preserved in clone 2, it also became more focal under increased selection (Extended Data Fig. 3 and Extended Data Fig. 1g). These analyses suggested that one pathway by which ecDNA structures evolve is through loss of non-selective (or disadvantageous) genes that were initially included in the amplicon. To examine this in single-molecule resolution, we employed combi-FISH, a method in which FISH probes specific for different amplicon regions that contain coding genes are used to determine their co-localization in ecDNA. Using a machine and deep learning approach (Extended Data Fig. 1h), we analyzed the co-localization of *DHFR* and *ARHGEF28* (found in 5q13.2), a gene originally included in the ecDNA of clone 3 due to a chromothriptic event^11^. Remarkably, although *ARHGEF28* frequently co-localizes with *DHFR* at low MTX pressure, it is almost completely lost at higher MTX concentrations (Fig. 1f-g and Extended Data Fig. 1i-j). The presence of multiple ecDNA species in single spreads at intermediate selective pressure, containing either *DHFR*, *ARHGEF28*, or both, is indicative of ongoing structural changes (Fig. 1g, see 800nM). Combi-FISH analysis of two genes initially found together with *DHFR* in the amplicon from clone 1 revealed that while *ACOT12* is mostly lost at 800nM (Extended Data Fig. 1k), *RASGRF2* is mostly still maintained at 10,000 nM resistance (Extended Data Fig. 1l). In line with this, reconstruction of ecDNA structures found in clone 1 using short-read^11^ and long-read DNA sequencing analysis^36^ (Fig. 1h), revealed significant structural heterogeneity. Although the closer proximity of *RASGRF2* to *DHFR* might explain why it was maintained at higher selective pressure, the functional impact of each gene might also play a role.

Strikingly, single cell RNA sequencing analysis revealed that while *DHFR* and *MSH3* are highly expressed in most cells, *ACOT12* and *RASGRF2* expression is mutually exclusive (Fig. 1i), revealing how ecDNA structural evolution drives functional divergence. Of relevance, genes found on ecDNA amplicons were among the most variably expressed (Extended Data Fig.1m) highlighting the numerical and structural plasticity of ecDNA. Such transcriptional variability was also observed in Colo320DM which harbors *MYC*+ ecDNA (Extended Data Fig. 1n), but not in HT-29 colorectal cancer cells that harbor a *MYC*+ HSR (Extended Data Fig. 1o), indicating that even in highly evolved cancer cell lines such as Colo320DM there are potential ongoing structural rearrangements in ecDNA that can generate heterogeneity (Fig. 1j and Extended Data Fig. 1n). Indeed, treatment of CHP-212 cells for two weeks with rapamycin (Fig. 1k) was sufficient to select for *MYCN*+ ecDNA structures that lost the previously reported rapamycin-sensitive passenger *DDX1* gene^37^ (Fig. 1l-m, Extended Data 1p-r), indicating that cancer cell adaptation could also be conveyed by structural evolution of ecDNA.

## Aggregation promotes ecDNA uneven inheritance

To study the mechanisms underlying ecDNA evolution, we examined how such acentric amplicons behave during mitosis. Using a computational imaging approach (Extended Data Fig. 4a), we measured ecDNA chromosomal tethering in mitotic *DHFR*+ ecDNA+ HeLa cells and observed a significant positive correlation (r=0.4746, p<0.0001) between ecDNA amount and reduced chromosomal tethering (Fig. 2a). ecDNA tethering was also significantly reduced (from ∼95% to ∼85% tethering) when transitioning from early (prophase-metaphase) to late (anaphase-telophase) mitosis (Fig. 2b-c and Extended Data Fig. 4b). A similar trend was observed in SNU-16 cells (Fig. 2d, Extended Data 4c), although loss of tethering was only significant for *FGFR2+* and not *MYC+* ecDNA, possibly because *FGFR2+* ecDNA are more abundant or due to a potentially different tethering mechanism.

**Figure 2:**
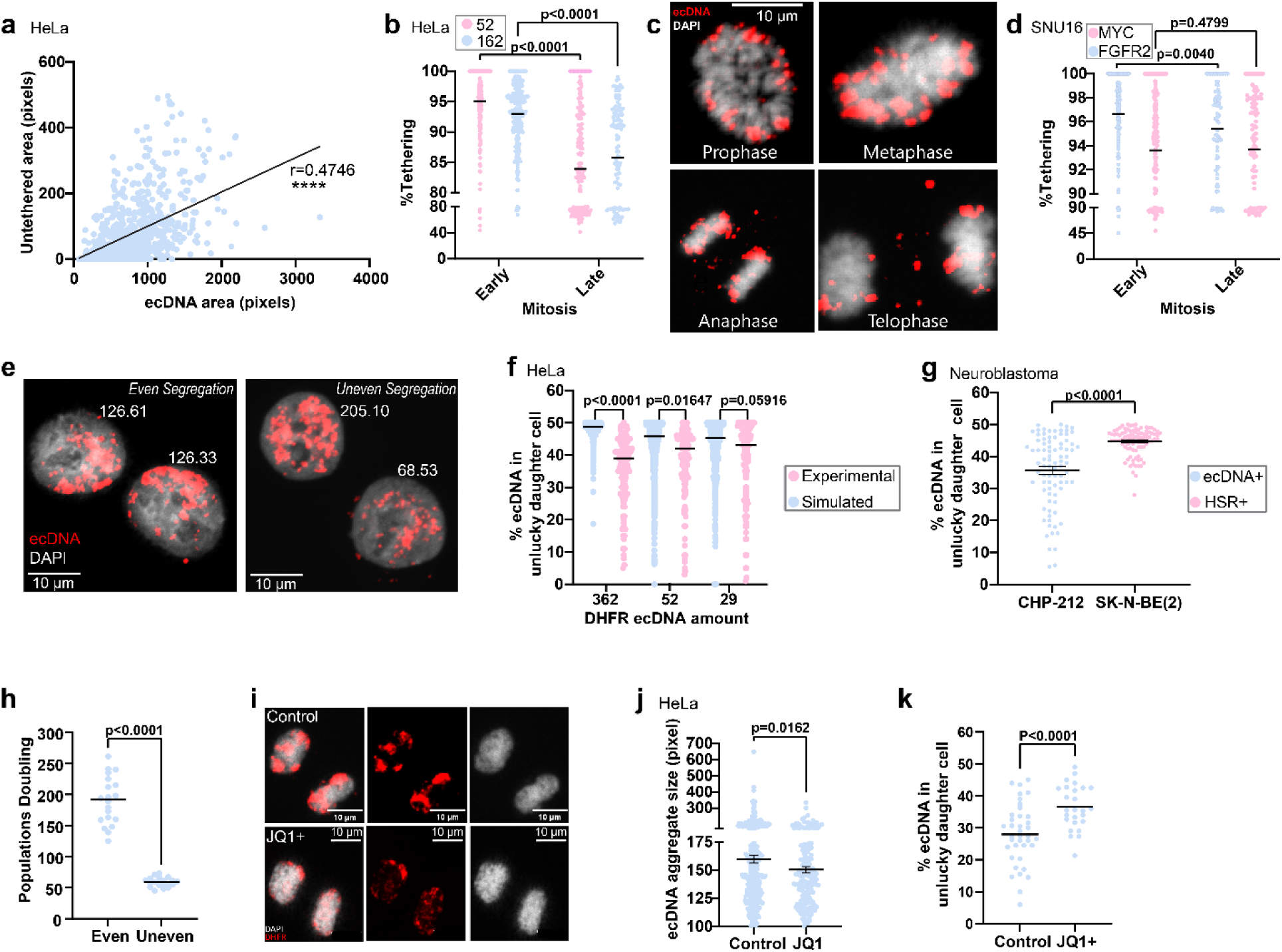
ecDNA aggregation drives uneven inheritance leading to rapid amplification. a, Correlation between ecDNA amount (ecDNA area, x axis) and untethered ecDNA (untethered area, y axis) as determined using DNA-FISH (see methods for computational details). Pearson correlation r=0.4746 one-tailed (n=3) shows a significant positive correlation between the overall amount of ecDNA and the amount of untethered ecDNA in mitotic cells. b, Quantification of ecDNA tethering during early and late mitosis in HeLa clone 1 cells with an average of 52 and 162 ecDNA. Two-way ANOVA followed by Šídák’s multiple comparisons test, mean ± SEM (n=3). c, Representative DNA-FISH images of ecDNA tethering in HeLa clone 1 cells with an average of 52 ecDNA. d, Quantification of *FGFR2*+ and *MYC*+ ecDNA tethering during early and late mitosis in SNU-16 cells. Linear mixed model fit, mean ± SEM (n=4). e, Representative DNA-FISH images of daughter HeLa clone 1 cells in interphase showing even and uneven segregation of *DHFR*+ ecDNA (red). Numbers denote ecDNA quantification (see methods for details). f, Quantification of ecDNA inheritance in HeLa clone 1 cells containing an average of 29, 52, and 362 ecDNA (pink). Y-axis presents the percent of ecDNA in the daughter cell inheriting the lower amount of ecDNA (“unlucky”). Simulations of ecDNA inheritance (blue) assuming random binomial distribution (p=0.5) for each ecDNA amount are presented. Comparison performed using pairwise nonparametric test. Mean±SEM (n=3) are presented. See method section for more details. g, Quantification of *MYCN+* ecDNA inheritance in CHP-212 cells (n=3). *MYCN*+ HSR+ SK-N-BE(2) cells were used as control (n=3). Welch’s t-test two-tailed; Mean±SEM. h, Simulation of the number of population doublings required for HeLa cells resistant to 1600nM MTX (containing an average of 74±80 ecDNA) to adapt to 3200nM MTX (reaching an average of 162±106 ecDNA). Comparison of even (p=0.5) and uneven (p=0.4, based on experimental data from Fig. 2f) is presented. Two-tailed student t-test. Mean±SD. i, Representative DNA-FISH images of HeLa clone 1 cells containing an average of 362 ecDNA treated with 500 nM JQ1+ or DMSO (control) for 24h. Note the reduced clustering on mitotic chromosomes in JQ1 treated cells. j, Quantification of ecDNA aggregate sizes tethered to mitotic chromosomes using DNA-FISH (as shown in panel i) in HeLa clone 1 containing an average of 362 ecDNA cell with or without JQ1 treatment. Size of larger (over 100 pixels) aggregates of tethered ecDNA is presented. Welch’s t-test (one tailed); mean+SEM (n=3). k, Quantification of ecDNA inheritance in CHP-212 cells treated with or without 500 nM JQ1 for 24h. Unpaired one-tailed student t-test (n=3). Mean±SEM.

Given that there are at least two possible fates for ecDNA in mitosis, chromosomal tethering or detachment, we sought to investigate the consequences of each fate. We assumed that chromosomally tethered ecDNA would be inherited in either of the two newly formed daughter cells following mitosis. We therefore measured the distribution of ecDNA in daughter cells following mitosis using interphase DNA-FISH (Fig. 2e-g). Surprisingly, we found that ecDNA is less evenly distributed in cells with higher levels of amplification (median distribution of 39% to one daughter cell and 61% to the other), which is opposite to simulated distributions rising from random segregation events (median distribution of 49% to one daughter cell and 51% to the other) for the same amount of ecDNA (Fig. 2e-f). A similar skewed distribution was observed in CHP-212 cells, but not in SK-N-BE(2) cells which harbor *MYCN*+ HSRs (Fig. 2g, Extended Data Fig. 4d). We reasoned that such a skewed distribution could facilitate accelerated accumulation of ecDNA copies over time and under selection. Indeed, comparing simulated random (p=0.5) and non-random (p=0.4, as determined by our experimental data) ecDNA mitotic distribution showed over three-fold decrease in the number of population doublings required to adjust from low to high selective pressure (Fig. 2h). This was consistent with our experimental data, in which cells undergoing a similar selective transition required an even shorter time to adapt to the higher MTX concentration (31±0.47 days, 5-fold difference from the simulated data). This faster adaptation could be explained by the fact that as cells divide, there is selection for increased ecDNA content, which in turn increases the rate of ecDNA accumulation (as more ecDNA results in less even distributions).

Increased gene amplification content should lead to higher transcription and subsequent protein production and therefore provide a potential selective benefit. Indeed, DHFR mRNA expression was positively correlated with DNA (R^2^=0.8784, Extended Data Fig.5a) and protein (Extended Data Fig. 5b-e) amounts, which consistently increased with higher MTX selection (Extended Data Fig. 5d,f-g), as previously reported ^38–40^. DNA-protein correlation of DHFR was also observed in single cells examined using IF-FISH analysis (Extended Data Fig. 5e). Similar correlations were found when examining *MYC*+ and *MYCN*+ ecDNA cell lines (Extended Data Fig. 5h-l), indicating that uneven ecDNA inheritance facilitates selective advantage across cancer types and amplified genes.

When examining mitotic figures of ecDNA+ cells, we frequently observed aggregates of ecDNA, reminiscent of ecDNA hubs^15^, tethered to chromosomes (Fig. 2i). Aggregation of ecDNA could essentially lower the number of dividing particles during mitosis, which could explain the increase in uneven distribution we observed in cells with higher ecDNA amounts. Indeed, we found that cells with more ecDNA contain larger aggregates, or hubs (Extended Data Fig. 4e). It was previously shown that treatment with the BRD4 inhibitor JQ1 disperses *MYC*+ ecDNA hubs^15^. Consistent with this, JQ1 treatment reduced *DHFR*+ ecDNA aggregates/hubs size in mitotic HeLa cells (Fig. 2i-j), while slightly increasing their overall amount (Extended Data Fig. 4f), potentially due to dispersal of large hubs into smaller ones. This resulted in a moderate increase in the evenness of ecDNA distribution in *DHFR*+ HeLa cells (Extended data Fig. 4g), and showed a stronger and more significant effect on *MYCN*+ CHP-212 cells (Fig. 2k), consistent with an essential role of ecDNA aggregation in skewed inheritance, and the previously observed co-segregation of distinct ecDNA species^2^. Importantly, these findings provide an additional explanation for why cancers often contain high amounts of ecDNA (beyond the need for higher protein production), as it drives increased aggregation and subsequent heterogeneity.

## ecDNA mis-segregate into distinct micronucleus types

Our analysis showed that most (>80%) ecDNA+ cells have one or more chromosomally detached ecDNA in mitosis (Extended Data Fig. 4b), raising the possibility that these will be frequently micronucleated. Indeed, live imaging of H2B-mCherry labeled ecDNA+ cells revealed that H2B+ minute particles frequently remain in the cytosol following mitosis (Fig. 3a and Supplementary Movie 1), as also observed in Colo320DM-GFP cells containing LacO-LacI-GFP labeled ecDNA^41^ (Extended Data Fig. 6a and Supplementary Movie 2). Examination of interphase nuclei using DNA-FISH revealed four classes of micronuclei in ecDNA+ cells (Fig. 3b): (1) DAPI+ only and (2) DAPI+/ecDNA+, both likely involve missegregation of a chromosome to which ecDNA are sometimes tethered or ‘hitchhiked’ (hereafter referred to as hitchhikers), (3) large ecDNA+, potentially originating from ecDNA aggregates/hubs (hereafter referred to as hubs), and (4) minute ‘nanonuclear’ ecDNA+, which represent a ∼single cytosolic ecDNA molecule. Types (3) and (4) represent a spectrum of ecDNA+ micronuclei sizes that failed to tether chromosomes during mitosis. While DAPI+ only micronuclei were more frequent in cells lacking ecDNA (6.22±1.14% in HeLa S3 and 6.87±0.86% HSR+ HeLa, 2.17-2.82(±1.85,1.7)% in ecDNA+ HeLa, Fig. 3c), the other micronucleus types were exclusively observed in ecDNA+ cells (ecDNA-negative cells showed residual amounts of signal), and their frequency significantly increased in HeLa cells containing more ecDNA (from 57±7% to 166±21% micronuclei in cells with an average of 52±65 and 362±323 ecDNA, respectively, Fig. 3c and Extended Data Fig. 6b), consistent with reduced tethering (Fig. 2a). High ecDNA+ micronuclei frequencies were also observed in CHP-212 (75±16%, Fig. 3d), Colo320DM (36±10%, Extended Data Fig. 6c), and SNU-16 cells (88±19%, Extended Data Fig. 6d), and in neuroblastoma patient samples (33±21%, Fig. 3e-f), but not in *MYCN*-SH-SY5Y or *MYCN*+ HSR+ SK-N-BE(2) cells (Fig. 3d) or patients without amplification (Fig. 3e). The predominant form of ecDNA+ micronuclei in all sample types appeared as a minute nanonuclear structure with an approximate size equal to the endogenous FISH signal, thus corresponding to a single ecDNA (referred to as “singlet” hereafter, Fig. 3f-i). Cells with more ecDNA (and with larger ecDNA aggregates, Extended Data Fig. 4e) had increased ecDNA+ micronuclei size (Fig. 3g).

**Figure 3:**
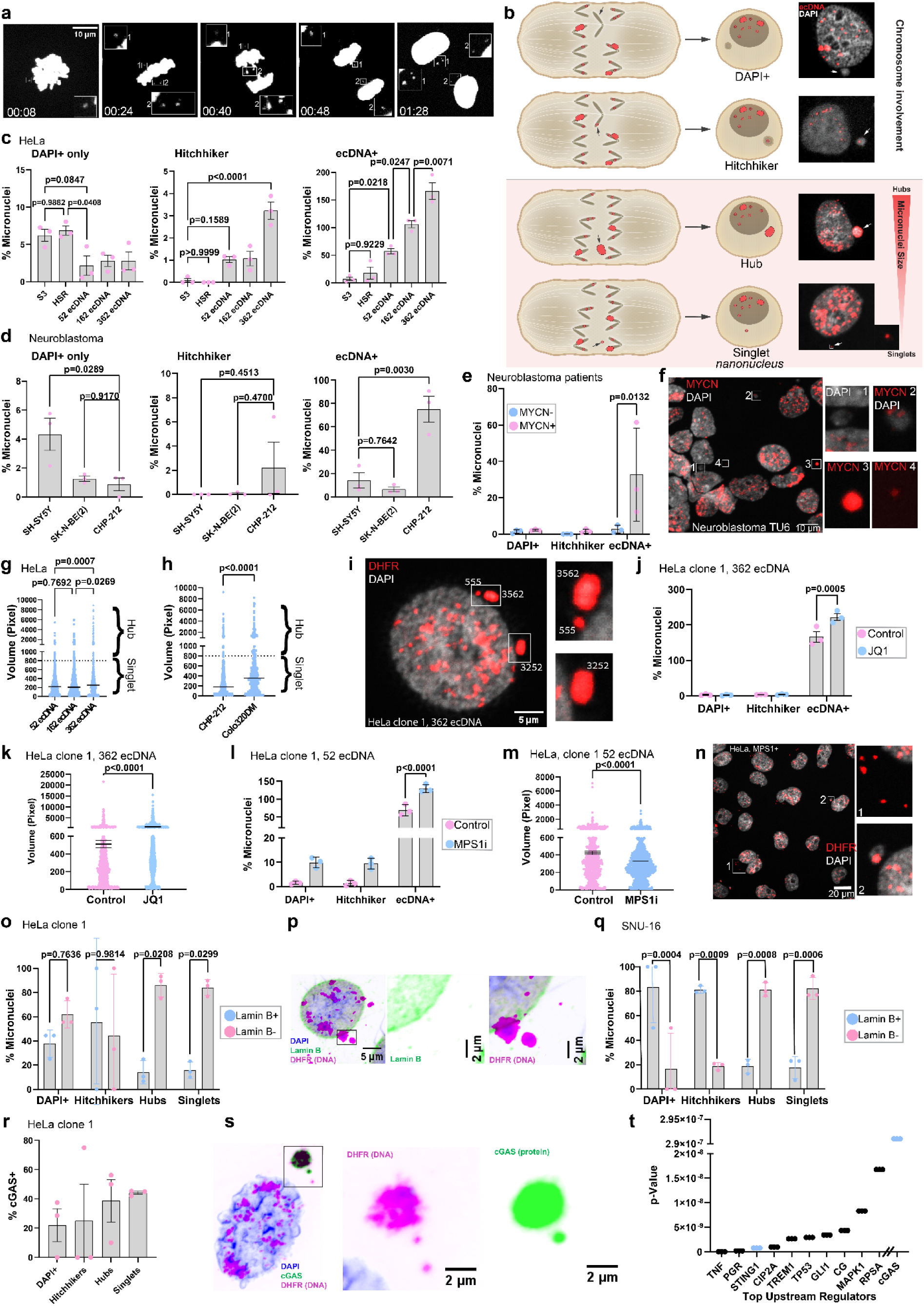
Mitotically detached ecDNA frequently entrap within distinct micronucleus types. a, Snapshots from Supplementary Video #1 showing a dividing H2B-mCherry+ HeLa clone 1 containing 362 ecDNA cell (from a population with an average of 362 ecDNA). Note the detached ecDNA during mitosis that remain in the cytoplasm following division. b, Description of the four micronucleus types and their origin as identified in this study. See main text for more details. c, Quantification of micronucleus types in HeLa cells without amplification, with HSR, and with increasing amounts of ecDNA. One-way ANOVA followed by Tukey’s multiple comparisons, mean ± SEM (n=3). d, Quantification of micronucleus types in neuroblastoma cell lines. One-way ANOVA followed by Tukey’s multiple comparisons test, mean ± SEM (n=3). SH-SY5Y – no amplification, SK-N-BE(2) – *MYCN*+ HSR, CHP-212 – *MYCN*+ ecDNA. e, Quantification of micronucleus types in six neuroblastoma patients (n=3 ecDNA-, n=3 ecDNA*+*). Two-way ANOVA followed by Šídák’s multiple comparisons test, mean ± SEM. f, Representative DNA-FISH image of *MYCN*+ ecDNA*+* neuroblastoma patient sample. Inserts show examples of DAPI+ (1), hitchhiker (2), hub (3), and singlet (4) micronucleus types. g-h, Quantification of ecDNA+ micronucleus sizes in HeLa clone 1 cells with an increased number of ecDNA (g) and in CHP-212 and COLO320DM cells (h). One-way ANOVA followed by Tukey’s multiple comparisons test, mean ± SEM (n=3). i, Representative DNA-FISH image of HeLa clone 1 cells with 362 ecDNA showing different sizes of ecDNA+ micronuclei. White numbers denote micronucleus size (see methods for details). j,l, Quantification of micronucleus types in HeLa clone 1 containing 362 ecDNA after 500 nM JQ1 treatment for 24h (j) or 500nM reversine treatment for 24h (l) treatment. Two-way ANOVA followed by Šídák’s multiple comparisons test, mean ± SEM (n=3). k,m, Quantification of ecDNA+ micronucleus sizes in HeLa clone 1 containing 362 ecDNA after 500 nM JQ1 treatment for 24h (k) or 500nM reversine treatment for 24h (m). Welch’s t-test two-tailed, mean ± SEM (n=3). n, Representative DNA-FISH image of HeLa clone 1 containing 52 ecDNA treated with 500 nM 24h reversine. o,q, Quantification of lamin B1+ micronuclei in a HeLa clone 1 containing 52 ecDNA (o) and SNU-16 cells (q). Two-way ANOVA followed by Tukey’s multiple comparisons test, mean ± SEM (n=3). p, Representative IF-FISH image of lamin B1 negative singlet and hub-type mirconuclei in HeLa clone 1 containing 52 ecDNA. r, Quantification of cGAS+ micronuclei in HeLa clone 1 containing 52 ecDNA. Data are shown as mean ± SEM (n=3). s, Representative IF-FISH image of cGAS+ singlet and hub-type micronuclei. t, Ingenuity pathway analysis showing top upstream regulators (top 10 and cGAS which was number 23) when comparing HeLa clones 1-3 at 10,000 nM MTX resistance to control HeLa S3 cells (n=4). Blue highlight the cGAS and STING pathways. Y axis shows p-value for each pathway.

Having established that untethered ecDNA frequently undergo micronucleation, we examined the contribution of DNA damage and repair, ecDNA hub dispersal, and elimination of the mitotic checkpoint to this process. As expected, treatment with etoposide, a type II topoisomerase inhibitor, significantly increased the amount of DAPI+ only (from 1.6±0.5% to 12.7±0.8%) and hitchhiker (from 1.4±0.9% to 24±3.8%) micronucleus types, however, with no apparent effect on ecDNA+ micronuclei or their size distribution (Extended Data Fig. 6e-f). Treatment with NU7441, a DNA-PKcs inhibitor, resulted in a moderate but significant increase in ecDNA+ micronuclei (p= 0.0205) (Extended Data Fig. 6g-h), indicating that ecDNA tethering might rely, in part, on intact non-homologous end-join (NHEJ) repair. Recent work found that treatment with JQ1 increases ecDNA micronucleation in *MYC* amplified cell lines but not in HeLa cells with low *DHFR* amplification (10-20 *DHFR*+ ecDNA, resistant to 80nM MTX^19^). Consistent with the effect we observed on ecDNA aggregation in HeLa cells (Fig. 2i-j), we detected a significant increase in *DHFR*+ hub-type micronuclei following JQ1 treatment (from 166±21% to 222±14%, Fig. 3j-k, Extended Data Fig. 6i), potentially due to incomplete hub dispersal leading to an overall increase in micronucleation. In contrast, inhibition of monopolar spindle 1 (MPS1) using reversine^42^ (MPS1i), significantly increased the formation of smaller, singlet-type ecDNA+ micronuclei (Fig. 3l-n), suggesting a possible role for premature exit from mitosis in ecDNA tethering.

To determine whether different micronuclear subtypes differ in structural integrity, we examined lamina properties across ecDNA+ and ecDNA-cells. Immunofluorescence combined with DNA-FISH (IF-FISH) revealed that micronuclei lacking chromosomal elements, namely hub- and singlet-type ecDNA+ micronuclei, were frequently lamin B- (∼80%), whereas hitchhiker and DAPI+ only micronuclei showed much higher lamin B staining. This pattern was consistent across multiple ecDNA+ models, including *DHFR*-amplified HeLa cells (Fig. 3o-p) and *MYC*/*FGFR2*-amplified SNU-16 cells, in which chromosomally-derived micronuclei were mostly lamin B+ (Fig. 3q). Lamin B deficiency was not attributable to micronuclear age (micronuclei persisting for several cell cycles), as induction of fresh micronuclei using MPS1i yielded no change in lamin B composition in either ecDNA- or ecDNA+ contexts (Extended Data Fig. 6j-m). Given that lamin B loss is associated with nuclear fragility, we hypothesized that ecDNA+ micronuclei are prone to rupture with consequent cytosolic DNA exposure and cGAS activation^43,44^. Indeed, cGAS immunostaining colocalized with >40% of hub and singlet micronuclei in *DHFR*+ HeLa cells and at similar levels in SNU-16 (Fig. 3r-s, Extended Data Fig. 7a). Transcriptomic profiling supported these findings: bulk RNA-seq of ecDNA+ HeLa cells revealed robust upregulation of cGAS, STING, and downstream interferon-stimulated genes, with scRNA-seq and proteomics confirming pathway activation across clones with increasing ecDNA copy number (Fig. 3t, Extended Data Fig. 7b-f). Collectively, these results demonstrate that ecDNA+ micronuclei are structurally fragile and frequently rupture, thereby leading to activation of the cGAS-STING signaling pathway in cGAS+ cells.

## Nanonuclear ecDNA degradation through autophagy

Previous studies showed that micronuclei can partially or completely degrade through autophagic processes^45–48^. The near-complete absence of lamin B and small size of ecDNA+ singlet nanonuclei prompted us to investigate if these might also undergo autophagic degradation, similar to the autophagy-mediated clearance of free genomic DNA that arises from collapsed micronuclei^49^. To test this, we treated ecDNA+ cells with the V-ATPase inhibitor bafilomycin A for 6-24 hours. Both LC3B and p62 levels robustly increased, indicative of efficient autophagic inhibition (Extended Data Fig. 8a-b), with some p62 puncti co-localizing with cytosolic ecDNA+ singlets (Extended Data Fig. 8b). Strikingly, singlet-type ecDNA+ nanonuclei significantly increased within 6 hours of treatment and reached ∼3-fold increase after 24 hours in *DHFR*+ ecDNA+ HeLa and *MYC*/*FGFR2*+ ecDNA+ SNU-16 cells (Fig. 4a-b, Extended Data Fig. 8c-d). Notably, bafilomycin A treatment did not affect the frequency of other micronucleus types (Fig. 4a and Extended Data Fig. 6c), nor did it affect ecDNA tethering (Extended Data Fig. 8e). To critically examine autophagic activity, we knocked down two essential components of the autophagy pathway: FAK family–interacting protein FIP200, which is part of early phagophore formation and the E2-like enzyme ATG3, which plays a role in autophagosome formation. Efficient knock-down of either protein (Extended Data Fig. 8f-g) resulted in a specific and significant increase (1.5-2-fold) of ecDNA+ singlets in both SNU-16 and HeLa cells (Fig. 4c-e) with no effect on ecDNA tethering (Extended Data Fig. 8h).

**Figure 4:**
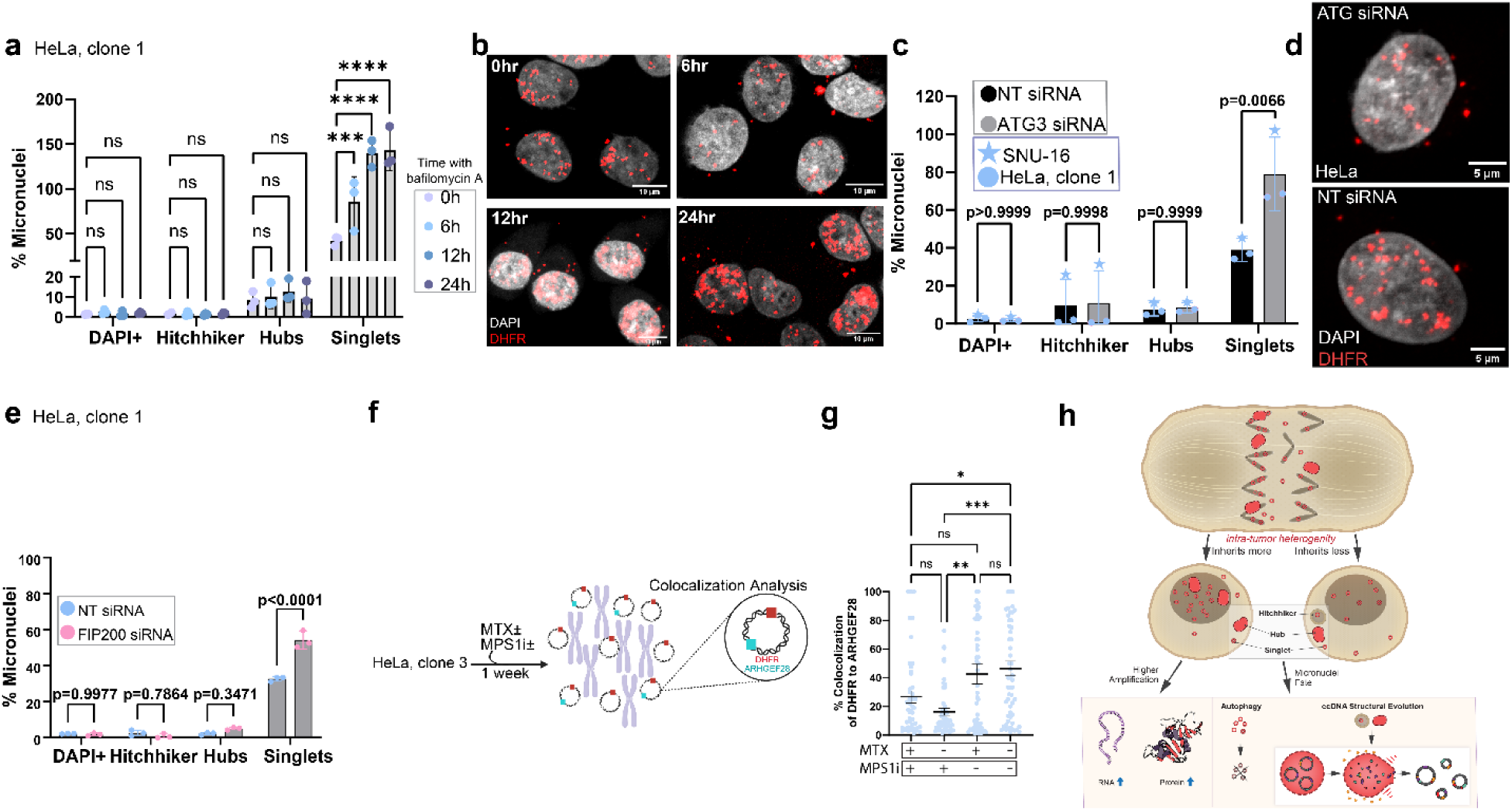
Autophagic degradation of ecDNA+ nanonuclei contributes to ecDNA structural evolution. a. Quantification of micronuclei percent (relative to primary nuclei) in a HeLa clone 1 containing 52 ecDNA following 100 nM bafilomycin A1 treatment for 6 to 24 hours. Two-way ANOVA followed by Tukey’s multiple comparisons test. Data are presented as mean ± SEM (n=3). *** p=0.0001, **** p<0.0001 b. Representative images showing of ecDNA+ micronuclei in HeLa clone 1 containing 52 ecDNA cells treated with 100 nM Bafilomycin A1 for 6 to 24 hours. c. Quantification of micronuclei percent (relative to primary nuclei) following ATG3 siRNA-mediated knockdown (72 hours) in HeLa clone 1 containing 52 ecDNA (n=2) and SNU-16 (n=1) cells. Two-way ANOVA followed by Tukey’s multiple comparisons test. Data are presented as mean ± SEM. d. Representative images showing increased DHFR⁺ ecDNA-containing micronuclei in the cytosol upon ATG3 knockdown of HeLa clone 1 containing 52 ecDNA. e. Quantification of micronuclei percent (relative to primary nuclei) following FIP200 siRNA-mediated knockdown (72 hours) in HeLa clone 1 containing 52 ecDNA. Two-way ANOVA followed by Tukey’s multiple comparisons test (n=3). f, Illustration of experimental set-up performed in panel g. HeLa clone 3 cells were treated with combinations of 200 nM reversine and 800 nM MTX for 1 week and metaphase spreads were prepared for DNA-FISH. g, Percent of colocalization of *DHFR* and *ARHGEF28* genes per cell in HeLa clone 3 treated with 200 nM ± reversine and 800 nM ±MTX (see panel f) for 1 week. One-way ANOVA with Dunnett’s multiple comparisons test (n=3). Data are presented as mean ± SEM. * p=0.0316, ** p=0.0012, *** p=0.0001 h, Graphical summary of study observations. ecDNA aggregation promotes uneven mitotic distribution leading to rapid accumulation and heterogeneity in the progeny. Cytosolic mis-segregated ecDNA encapsulate in different micronucleus types, driving structural evolution through potential chromothriptic rearrangements or autophagic degradation.

The evolution of ecDNA structures, including the selection of ecDNA species with different genetic content (Fig. 1f-g), is thought to occur via ongoing structural rearrangements^11^, but could also be directed by ecDNA micronucleation and degradation. To test this possibility, we treated ecDNA+ HeLa cells with reversine for one week (Fig. 4f), which increased ecDNA+ micronuclei by 1.8-fold and hitchhikers by ∼8-fold (Extended Data Fig. 8i), and examined its effect on the loss of *ARHGEF28* (which is lost under increased MTX selection, Fig. 1f-g). Combi-FISH analysis revealed a ∼3-fold significant reduction in the co-localization of *DHFR* with *ARHGEF28*, following reversine treatment with or without MTX (Fig. 4g), suggesting that excessive formation of ecDNA+ nanonuclei and consequent degradation accelerates the evolution of ecDNA. Overall, these data suggest that ecDNA micronucleation or degradation shape the evolutionary trajectories of ecDNA, thereby contributing to cancer cell adaptation.

## Discussion

Our findings reveal that rapid ecDNA accumulation across cell generations is driven by aggregation, which also fuels intra-tumoral heterogeneity (Fig. 4h). This dynamic mode of inheritance endows cancers with increased plasticity and accelerates their evolutionary potential, helping explain the strong selective advantage conferred by ecDNA^5^. By linking ecDNA amplification to micronucleation, this study unites two fundamental hallmarks of cancer genome evolution. While previous work highlighted the role of micronuclei in the genesis of ecDNA through chromothripsis^11,50^, our results establish ecDNA micronucleation itself as a key evolutionary process—one that not only facilitates structural rearrangements but also enables active clearance of ecDNA (Fig. 4h), thereby shaping the adaptive landscape of the cancer genome.

Beyond their structural role, ecDNA-containing micronuclei exhibit functional consequences with broad implications for cancer progression. Their fragility and frequent rupture trigger innate immune signaling through the cGAS–STING pathway, suggesting a mechanism by which ecDNA may impact metastatic potential^7,51^. Importantly, we show that distinct classes of micronuclei follow different fates, raising the possibility that shifting the balance among them could be therapeutically exploited. For example, targeted strategies that drive singlet-type ecDNA micronuclei toward autophagic degradation could reduce ecDNA burden and blunt tumor adaptability. In this view, ecDNA inheritance is not merely a mechanism of gene amplification but a dynamic process that dictates structural fate, cellular plasticity, and ultimately cancer evolution, while simultaneously offering new avenues for intervention.

## Methods

### Cell culture

Human tumor cell lines were obtained from the American Type Culture Collection (ATCC) or were created in our lab. HeLa cells were grown in Dulbecco’s modified Eagle’s medium (DMEM, Thermo Fisher Scientific; 41965-039) containing 10% fetal bovine serum (FBS, Thermo-Fischer; A5256701) supplemented with 100 U/ml penicillin (Diagnovum; D910-100ML), 100 U/ml streptomycin (P/S, Diagnovum; D910-100ML), and 2 mM L-glutamine (Biological Industries; 03-020-1B), at 37 °C under 5% CO_2_. When treating with methotrexate (MTX, calbiochem; 454126-100MG), dialyzed FBS was used instead (Hyclone; SH30079.03 Cytiva) The cells were split twice a week, and when grown with MTX, treated twice a week. In the isogenic system (clone 1-PD29429h, clone 2-PD29425d, clone 3-PD29427k), cells started out with 40 nM MTX resistance. Resistance of higher level of MTX was created by seeding 200,000 cells in a 6 well plate and treated with the wanted dose of MTX, which usually was double the dosage of MTX from the previous one. Cells that were undergoing the resistance challenge were maintained until they maintained confluency (minimum of two weeks) in the 6 well plate. SH-SY5Y/CHP-212/SK-N-BE(2) cell lines were grown in DMEM F12 (Thermo fisher Scientific; 21331020), 10% FBS, 1% P/S, and 2 mM L-glutamine. IMR-32 were grown in EMEM F12 (Sigma; M5650-500ML). Colo320DM, Colo320DM-GFP, UACC-1594, NCI-H524, SNU-16 and DLD1 cell-line were grown in RPMI-1640 (Gibco; 21875-034) 10% FBS, 1% P/S, and 2 mM L-glutamine. All cells underwent routine mycoplasma testing and were found negative.

### Drug Treatments

Chronic MPS1 Inhibitor: 200,000 HeLa cells from clone 3 were seeded in a 6-well plate, or 50,000 cells on a coverslip. 24h later medium was changed and 200 nM Reversine (Sigma; R3904-1MG) was added. DMSO concentration did not exceed 0.001%. After three days cells either had their medium changed or split into media with fresh drugs. After a week the cells underwent medium change with no drugs and a 15 hours later were pelleted or received 100 ng/ml colcemid (KaryoMax, Gibco; 5212012) and metaphase spreads were created.

Rapamycin treatment: 200,000 CHP-212 cells were seeded in a 6-well plate. 24 hours later, the cells were treated with 1µM of Rapamycin for a period of 13 days. On day 13, the cells were split and on day 16 the cells underwent fixation with 3:1 methanol: acetic acid and subjected to DNA-FISH.

Acute BRD4 inhibitor: For shake-off experiment: A confluent 10 cm plate of HeLa clone 1 resistant to 10,000 nM MTX received a treatment of 500nM of JQ1 (Sigma; SML1524-5MG) for 24 hours. Cells were then shook off to enrich for mitotic cells. 20,000 mitotic cells were seeded on a poly-lysine coated coverslip with new medium. 24 hours later the plate underwent fixation with 3:1 methanol: acetic acid and subjected to DNA-FISH. For coverslips: 50,000 cells were seeded on poly-lysine coated coverslips and medium. After 4 hours cells received 500 nM of JQ1 for 24 hours. After 24 hours the coverslips were fixed with 3:1 methanol: acetic acid and processed for DNA-FISH.

Acute drug treatments: HeLa clone 1 cells were seeded on coverslips and 24 h afterwards had their medium change with one of the following drugs: Reversine (500 nM, 24h), JQ1 (500nM, 24h), Nu7441(Selleckchem; s2638, 10 µM, 48 h), Etoposide (Sigma, E1383; 5µM for two hours and release for 24 hours). Afterwards the coverslips underwent fixation with 3:1 methanol: acetic acid and subjected to DNA-FISH.

Fluorescent Methotrexate: HeLa clone 1 cells were detoxed from normal methotrexate for a week before receiving 10µM of fluorescent methotrexate (Biotium; 00023) for 24h. Cells were sorted using SORP FACSAria3 by the Weizmann life science core facility. Gating was done using an HSR *DHFR*+ clone. Bafilomycin Treatment: Glass coverslips were pre-coated with poly-L-lysine (Sigma-Aldrich;P4707) for 5 mins at room temperature, rinsed with sterile water, and allowed to dry completely. HeLa clone 1 or SNU-16 cells were seeded onto sterile glass coverslips placed in 24-well culture plates and allowed them to adhere overnight under standard growth conditions. The following day, cells were treated with Bafilomycin A1 (Sigma-Aldrich; B1793) at a final concentration of 100 nM diluted in complete culture medium. Treatment durations were 0, 6, 12, and 24 h. For each time point, cells were maintained at 37°C in a humidified 5% CO₂ incubator. Control samples (0-hour) received an equivalent volume of vehicle (DMSO). At the end of each treatment time point, cells were washed once with 1× PBS and fixed directly on coverslips with 100% Methanol at −20°C for at least 20 minutes.

### siRNA-mediated Knockdown of ATG3 and FIP200

For targeted knockdown of *ATG3* and *FIP200*, HeLa clone 1 or SNU-16 cells were transfected with a DharmaFECT 1 (T-2001-03) according to the manufacturer’s instructions. Cells were seeded at appropriate densities to ensure 70% confluency at the time of transfection. For *ATG3* knockdown, the following siRNAs were used as a pool: OTP si-ATG3 #1: Dharmacon, Cat. No. J-015375-05, OTP si-ATG3 #2: Dharmacon, Cat. No. J-015375-06, OTP si-ATG3 #3: Dharmacon, Cat. No. J-015375-07, OTP si-ATG3 #4: Dharmacon, Cat. No. J-015375-08.

Cells were transfected with siRNA, and the medium was replaced 24 hours post-transfection. Cells were then incubated for a total of 72 hours from the time of transfection. At 48 hours post-transfection, cells were trypsinized, counted, and re-seeded onto poly-L-lysine–coated coverslips for FISH analysis, and in parallel onto separate dishes for Western blotting to assess knockdown efficiency.

For *FIP200* knockdown, siRNA specific to human FIP200 (RB1CC1, Cat# EHU043821 Sigma-Aldrich) was used and transfected and processed as described above.

Knockdown efficiency was confirmed at the protein level by Western blot using antibodies specific to ATG3 (Sigma Aldrich, A3231 at 1:1000 dilution) and FIP200 (CST, #12436). Downstream analyses, including DNA-FISH was performed on fixed cells at the 72-hour time point as described in the corresponding methods sections.

### DNA-FISH

Chromosome spreads were obtained by dropping mitotic cells onto glass slides. In brief, cells were arrested for 4 h with 100 ng/ml colcemid (KaryoMAX, Gibco; 15212012) and then incubated in 75mM KCl for 10 min at 37 °C. Cells were fixed by adding methanol/acetic acid (3:1, ACS grade or above), washed with fixative three times, and kept in fixative at −20 °C until use. Coverslips were fixed in 1 mL methanol/acetic acid (3:1) for a minimum of 20 minutes and then air-dried. Genomic DNA and probes were applied and co-denatured at 75 °C for 5 min by placing slides on a pre-heated metal plate. Samples were hybridized overnight at 37 °C in a dark humidified chamber. Slides were subsequently washed with 0.4 × SSC at 72 °C for 2 min and rinsed in 2 × SSC, 0.05% Tween-20 at room temperature for 30 s. Slides were then rinsed in Milli-Q water, counterstained with DAPI (Sigma; 10236276001) and mounted using pro-long gold (Thermo-Fischer; P36935). FISH images were acquired on Nikon TI2 confocal spinning disc Yokogawa at 60× magnification. Maximum intensity projections were generated using the ImageJ program. Amplification was manually counted, with a minimum of 30 chromosome spreads per sample.

### IF-FISH and analysis

Cells were seeded on coverslips and fixed with Methanol for 20 minutes in -20 °C. DNA-FISH was performed, and then coverslips were blocked with Triton Blocking buffer for 1h in RT. Coverslips were next incubated in 4 °C with a 1^st^ antibody (Supplementary materials). Coverslips were washed with PBST three times, 5 min each time, and incubated with a secondary fluorescent antibody (Abcam, ab150077, 1:500 or Abcam, ab150113 1:500) in RT for 1h. Coverslips were subsequently washed with PBST three times, 5 min each time. Coverslips were then counterstained with DAPI and mounted using pro-long glass (Thermo-Fischer; P36980) onto slides.

Manual scoring of Lamin-B1 and cGAS signals was performed on micronuclei (MN) identified based on DAPI and FISH fluorescence. For each experimental condition, at least 100 micronuclei were analyzed across multiple randomly selected fields of view. Only well-preserved cells with intact morphology and micronuclei that were in focus were included for quantification.

Lamin-B1 Positivity Criteria: Micronuclei were considered lamin-B1 positive if they exhibited either a continuous or partial lamin-B1 rim signal encircling the MN (applicable to hitchhiker and negative-type MN), or a clear colocalized lamin-B1 fluorescence overlapping with the FISH signal in singlet and hub-type MN.

cGAS Positivity Criteria: cGAS localization was scored as positive when cytosolic cGAS signal clearly accumulated and overlapped with the DAPI or FISH-positive micronuclear structures. All image processing and quantification were performed using FIJI (ImageJ).

### TISSUE-FISH

TU (tumor) preparation is mechanically disrupting solid tumor from fresh tissue smeared on slide with Methanol and Glacial Acetic Acid. In a Phase-contrast microscopy (PCM) probe addition (MYCN, MetaSystems. ON-D-6031-100-OG) and hybridization areas were chosen. Coverslip was added after air-drying and denatured at 75 °C for 2 min by placing slides on ThermoBrite. Hybridization was at 37°C for 24 h. The slide was washed in 0.4 × SSC/0.3%NP-40 (PH7.5) at 73 °C for 2 min, followed by 2 × SSC/0.1%NP-40 (PH7.0) at RT for 1 min. The nuclei were counterstained with DAPI. Nuclei with two visible signals were scored as negative for *MYCN* amplification. According to the guidelines, the cutoff point for *MYCN* amplification was set at 10 signals (amplification ≥10; gain <10).

### Microscopy

Samples were imaged using Spinning Disk confocal microscope (Nikon Instruments Inc., Melville, NY) with 50µm pinhole (Yokogawa CSU-W1). Images were acquired with a CFI Plan Apochromat IR X60/1.27 objective, with GFP, RFP, CY and DAPI filters with sCMOS camera (Photometrics, PRIME –BSI). Images were collected in Z-stack mode with 0.2 µm steps. The microscope used NIS-Elements AR 5.42.01 program. Live cell imaging was performed at 37°C under 5% CO_2_. Z-stacks of 25 sections at 1-μm intervals were acquired every 8minutes for 16 hours.

### Simulations of ecDNA segregations

For the single-generation simulation, actual counts from each MTX (100, 800 or 10000) of HeLa clone 1 were used as the starting point. Each ecDNA count was split randomly (binomial distribution with a probability of 0.5) to two daughter cells. The percentage (from the parent total) was calculated for each daughter cell. This process was repeated 1000 times to create a distribution of possible values. The simulated percentage values under 50% were compared to the real experimental values using Mann-Whitney tests.

For the multiple-generation simulation, the starting point was a random population of 2000 cells with a normal distribution (mean=70, s.d.=20). In each generation, each cell was split randomly (binomial distribution with a probability of 0.5 or 0.4 for symmetric and skewed simulations, respectively).Cells with less than 10 counts were removed in each step, and the values in each daughter cell were multiplied by 2 before the next generation. We kept running more generations until the mean ecDNA counts reached 160. We repeated this whole process 20 times, and noted the number of generations it took for the mean value to increase from 70 to 160. We compared the number of required generations between the symmetric and the skewed simulations with a t-test.

All statistics and simulations were run using R, v. 4.5.0.

### Probe Preparation for DNA-FISH

To make custom BAC probe: BACs (see table in supplementary data) were grown in single-clones with 12.5 μg/mL chloramphenicol in 250 mL bacterial cultures. DNA was extracted using NucleoBand Xtra BAC Large contract DNA purification kit (Macherey-Nagal; MAN-740436.25). Isolated BACs underwent nick translation using Enzo Nick Translation DNA Labeling System 2.0 (ENZ-GEN111-0050) together with the following labeled dUTP: Orange 552 dUTP (Enzo; ENZ-42842), Green 448 dUTP (DYOMICS; DY-448-34). Labeled BAC probes were suspended in hybridization buffer (50% ultra-pure deionized formamide, 0.05% (v/v) 1 M NaPi, 0.1% (v/v) 20× SSC, 10% dextran sulfate), finalized with Milli-Q water and adjusted to pH 7.0.

### Taq-Man

1,500,000 cells were counted, harvested, and made into pellets. DNA was purified (QIAGEN DNeasy Blood and Tissue Kit, 69504) and measured for purity and concentration using NanoDrop 8000 (Thermo Scientific, 010015885). For calculations:1.5x10^6^ RPE1 cells have 3x10^6^ MYCN copies in 100 µL. Six 1:5 serial dilutions were prepared. Samples diluted 1:100 and run in triplicates. PCR reaction mix was prepared per TaqMan protocol with no adaptations (ThermoFisher. MasterMix, AB-A30866 3, MTO for DHFR and MYCN, AB-4400291). Plate was run on QuantStudio 5. Results were analyzed using a standard curve with the diploid RPE-1. Analysis was run on QuantStudio Real-Time PCR software. The three technical repeats were averaged into one value for 1-way ANOVA analysis, and copy number was normalized using nanodrop concentrations.

### Western blot

#### For MYC/MYCN/DHFR

Cells were harvested using Trypsin B. 1,000,000 cells were lysed using a mixture of 50 μL of RIPA buffer (Sigma; R0278) and a protease inhibitor (Sigma; P8340) for 20 min, in which every 5 min, the cells undergo a 5 s vortex. After 20 min, cell lysates underwent a 20 min centrifuge at 4 °C at 18,213 rcf. The lysates were then separated from the cell membrane remains. Protein concentration was determined by the BCA analysis, and 20 μg of protein was taken from each sample, supplemented with 4× loading buffer that has 0.02 mM DTT, and heated for 10 min at 70 °C. Samples were loaded into a 4–20% mPAGE bis-tris gel (Merck; MP42G12) at 140 V for 70 min and then transferred into nitrocellulose membrane (Biorad; 1704158) using Trans-Blot Turbo transfer system (Biorad; 1704158). The membrane was stained with Ponceau (Sigma; 6226-79-5) to assure the quality of the transfer. The membrane was de-stained with ddH2O and then blocked for 60 min with a blocking buffer (5% BSA in TBS-T). Next, the membrane was incubated overnight at 4 °C with primary antibody (supplementary materials). The next day, the membrane was washed three times for 10 min with TBS-T. The membrane was incubated with anti-rabbit/mouse IgG, HRP-linked secondary antibody (CST) for 1 h at room temperature, and washed four times for 7 min each with TBS-T. The membrane was imaged using ChemiDoc XRS+ (BioRad).

#### For LC3B, P62, ATG3, FIP200

Total cellular protein extracts were prepared in RIPA buffer (0.1 M NaCl; Bio-Lab Ltd, 21955 5 mM EDTA (J.T. Baker; 8993), 0.1 M sodium phosphate, pH 7.5; Sigma, 342483, 1% Triton X-100 (Sigma; X100), 0.5% sodium deoxycholate (Sigma; D6750), 0.1% sodium dodecyl sulfate (Sigma; L4509)) with a protease inhibitor cocktail (PIC; Merck, 539134). The extracts were centrifuged at 13000 rpm for 15 min at 4°C, and protein concentrations were determined using Thermo Scientific Pierce BCA Protein Assay Kit (Thermo Scientific; PI23227). Total proteins (30 µg) were separated by SDS−PAGE (Novex WedgeWell 8 to 16%, Tris-Glycine, Mini Protein gel (Invitrogen; XP08165BOX) for Fip200, p62, and ATG3) or Novex WedgeWell 4 to 20%, Tris-Glycine, Mini Protein gel (Invitrogen; XP04205BOX) for LC3B and transferred to a nitrocellulose membrane (Bio-Rad; 1704159). The membrane was blocked in PBS with 5% skim milk for 1 h at room temperature, and then incubated with the appropriate primary antibody overnight at 4°C. It was then washed three times with PBS-T (Tween-20 0.1%, Sigma; P1379) and incubated with the secondary antibody (goat anti-mouse or goat anti-rabbit) for 1 h at room temperature. Finally, the membrane was washed three times and specific proteins were visualized using the Enhanced Chemiluminescence (ECL) detection system (Biological Industries; 20-500-120).

### Computational scripts

#### Shakeoff and intensity within interphase cells

To measure the amplification intensities and areas within nuclei, nuclei were segmented using Cellpose 3.0 with the pretrained cyto3 model^52^. Z slices of the amplification were averaged and DAPI signals were segmented via pixel classification using the Trainable WEKA Segmentation plugin in FIJI^53^.

#### Micronuclei quantification

Nuclei were segmented using Cellpose-SAM^54^ and were counted using FIJI^53^. Micronuclei were segmented via pixel classification using Arivis Pro software (arivis AG, Rostok, Germany), followed by the exclusion of nuclear objects. Areas and intensities of the segmentations were calculated using Arivis Pro.

#### Combi-FISH

3D analysis was carried out to quantify the colocalization between the *DHFR* and *ARHGEF28* in HeLa clone 3 and between *MYCN* and *DDX1* in CHP-212 within spread nuclei using Arivis software (Carl Zeiss Microscopy GmbH, version 4.2.1). Spread nuclei were segmented using deep learning^55^. RFP and Cy5 were segmented using machine learning and were colocalized at distances lower than 500nm between the two geometries, with no consideration to the intensity. GFP was segmented using blob finder and was considered colocalized with RFP or Cy5 when there was overlap.

#### Image and Computer Vision Analyses^56^

Tethering of ecDNA to mitotic chromosomes was measured in two cell line populations: HeLa clone 1 (with increasing amount of ecDNA) and SNU-16, with targeted analyses to each of them. DAPI and ecDNA channels were measured. First step was to segment the signals in the different channels. Different algorithms applied to segment the DAPI and ecDNA signals due to the different signal characteristics. DAPI signal was subjected to Gaussian Blurring with a kernel size of 3x3, applying the OpenCV implementation, cv2.GaussianBlur. Then a fuzzy c-means clustering algorithm was applied^57,58^. A threshold value was selected to binarized the image and return the segmented signal in a binarized form. ecDNA segmentation: To segment the ecDNA signal, first, the OpenCV Adaptive Mean Thresholding (cv2.adaptiveThreshold()) implementation was applied to obtain the binarized segmented signal. Then, small objects under 10 pixels were removed. This analysis measured particle size on and outside DAPI segmentation.

HeLa clone 1 analysis: Three features were calculated in those analyses. The first one, F1, is the number of pixels in the binarized ecDNA signal which is given by the summation of the pixels. The second feature, F2, is the number of pixels of the binarized ecDNA signal which is completely not overlapping with the binarized DAPI signal. F2 was initialized to zero. Then, the Connected Components Analysis of OpenCV implementation (cv2.connectedComponents()) was applied to label all of the ecDNA. Next, the ecDNA labels were iterated and each of them was binarized. Next, the overlap with the binarized DAPI signal was calculated. If there was not any overlap for a given binarized ecDNA, then its pixels sum was calculated and added to F2. The third feature, F3, is a list that contains the number of pixels of each ecDNA foci that has at least partial overlap with the binarized DAPI signal. Here, when iterating over the ecDNA labels, the overlap between a given ecDNA and DAPI binarized signals was calculated and if there was at least partial overlap then the number of pixels of the given ecDNA was calculated and appended to the list to construct F3.

SNU-16 analysis: In that analysis, features F1-F2 as in HeLa were calculated for the MYC gene and FGFR2 gene with DAPI. This yields in total four features, F1-F4. Furthermore, three additional features were calculated, F5-F7. In F5, Connected Components Analysis was applied to both, the binarized MYC and FGFR2 signals in each image. This enables calculating for all the binarized ecDNA of MYC and FGFR2 signals the overlap with DAPI as in F3 in previous section. For those that are not in overlap with the binarized DAPI signal, the number of overlaps between the binarized MYC and FGFR2 ecDNA was calculated to construct F5. F6 and F7 are the number of binarized ecDNA of MYC/FGFR2 that do not overlap with the binarized DAPI and in overlap with the other binarized ecDNA signal, FGFR2/MYC, respectively.

Code: All code for those analyses was developed in Python 3.10.4. Specifically, OpenCV library (v.4.11.0) (https://opencv.org/) fuzzy c-means module (v.1.7.2)^57^, and scikit-image library (v.0.23.2)^58^ were utilized for the image and computer vision analyses. Additionally, NumPy library (v.1.26.4)^59^ was applied for array manipulations and Matplotlib library (v.3.9.0)^60^ for visualization.

### Whole Genome Sequencing Analysis

#### Illumina data processing

For HeLa clone 1 PD29429i raw paired-end whole-genome sequenced Illumina reads were subjected to a quality control (QC) assessment using *FastQC (Andrews, 2010)* v0.12.1. Adapter trimming and quality filtering were performed with *Trim Galore!* (https://github.com/FelixKrueger/TrimGalore, parameters: -q 20 --phred33 --illumina --paired -j 8 --length 20). Reads passing QC were aligned to the reference genome *GRCh38.p7* using *BWA-MEM* (Li H. and Durbin R. 2009) 0.7.17 with default parameters. PCR duplicates were subsequently identified and removed with *Picard* (http://broadinstitute.github.io/picard/) 3.1.1. To generate coverage profiles, the deduplicated BAM files were processed with *deepTools bamCoverage* (*Ramırez et al. 2016)* 3.5.1 using standard parameters. Structural variant (SV) calling was performed using *LUMPY* 0.2.13, and genotyping of identified variants was carried out with *svtyper* 0.7.1, both run with standard parameters against the *GRCh38.p7* reference genome assembly.

#### Nanopore sequencing and data processing

Libraries were prepared with the ONT Ligation Sequencing Kit (LSK-109) together with the Native Barcoding Expansion kit (NBD104) to enable multiplexing. A maximum of four samples were multiplexed per sequencing run, starting from 248 ng of input DNA per library. Nanopore whole-genome sequencing was performed on R9.4.1 MinION flowcell (FLO-MIN106; Oxford Nanopore Technologies). Sequencing was carried out according to the manufacturer’s standard protocols.

The base-calling was performed using *Guppy* v5.0.14 (guppy_basecaller, dna_r9.4.1_450bps_hac model). The obtained barcoded reads were demultiplexed using guppy_barcoder. Reads were then quality-filtered using *NanoFilt (De Coster et al. 2018)* 2.8.0 (-l 100 --headcrop 50 --tailcrop 50) and aligned against the human reference genome (*GRCh38*) using *ngmlr* (Sedlazeck et al. 2018, in this paper both ngmlr and sniffles were published) 0.2.7. Structural variants (SVs) were identified with *Sniffles (Sedlazeck et al. 2018)* 1.0.12 with parameters --min_homo_af 0.7 --min_het_af 0.1 --min_length 50 --min_support 4. To obtain coverage profiles, the resulting BAM files were processed with *deepTools bamCoverage (Ramırez et al. 2016)* 3.5.1 using default parameters. The pipeline is available under https://github.com/henssen-lab/nano-wgs.

#### Decoil reconstruction

Reconstruction of ecDNA was performed using *Decoil* 2.0.0a1 [https://github.com/madagiurgiu25/decoil-pre, commit c4e10f4] in the consensus mode, developed as extension of the original *Decoil*^36^ implementation for this analysis to support short-read sequencing data and custom multi-VCF structural variant (SV) files. To achieve higher-resolution reconstructions, we applied *Decoil* in **consensus mode**. Consensus SV call sets were generated by selecting all unfiltered variants calls shared between nanopore- and Illumina-based per sample. The merging of SV calls was performed with *SURVIVOR* 1.0.7, generating multi-VCF files that served as input for *Decoil*. In this mode, reconstruction was performed using Illumina coverage profiles and the parameters: --fragment-min-cov 10 --fragment-min-size 50 --fragment-max-cov 2000 --min-vaf 0.1 --min-cov 10 --filter-score 20 --sv-caller multivcf. The reconstruction threads were visualized using *Decoil-viz*^36^ 1.0.4 (https://github.com/madagiurgiu25/decoil-viz) using the reference genome GRCh38/hg38 and annotation GENCODE V42.

#### CNV analysis

Copy-number analysis of HeLa clone 1 sequenced by whole-genome Illumina sequencing, was performed using *CNVkit* 0.9.10^61^ with the hmm-tumor segmentation method and default parameters.

### Bulk-RNA-Seq

Cells from 6-well plate were pelleted and flash frozen in liquid nitrogen. RNA was extracted using RNeasy Mini Kit (Qiagen; 74104), was checked for purity using Agarose gel, Nanodrop and Tapestation (all with a score of 9.8 and above). RNA amount was quantified using Qubit RNA BR Assay Kit (Thermo Fischer Scientific; Q10211).

RNA-seq libraries were prepared at the Crown Genomics Institute of the Nancy and Stephen Grand Israel National Center for Personalized Medicine, Weizmann Institute of Science. A bulk adaptation of the MARS-Seq protocol^62,63^ was used to generate RNA-Seq libraries for expression profiling of the cell-lines. Briefly, (30 ng of input) RNA from each sample was barcoded during reverse transcription and pooled. Following Agencourct Ampure XP beads cleanup (Beckman Coulter), the pooled samples underwent second-strand synthesis and were linearly amplified by T7 in vitro transcription. The resulting RNA was fragmented and converted into a sequencing-ready library by tagging the samples with Illumina sequences during ligation, RT, and PCR. Libraries were quantified by Qubit and TapeStation as well as by qPCR for the *GAPDH* gene as previously described^62,63^. Sequencing was done on a NovaSeqX plus using a kit of 1.5B 100 cycles, allocating 1600M reads in total (Illumina).

Single end reads with the median depth of 12,204,124 reads per sample were analyzed using the bioinformatics analysis pipeline User Friendly Transcriptomic Analysis Pipeline (UTAP)^64^. Reads were trimmed using cutadapt^65^ v4.1 (parameters: -a AGATCGGAAGAGCACACGTCTGAACTCCAGTCAC -a“A{10}” –times 2 -u 3 -u -3 -q 20 -m 25). Reads were mapped to the human genome hg38 using STAR^66^ v2.7.10a (parameters: –alignEndsType EndToEnd,–out Filter Mismatch Nover Lmax 0.05,– two pass Mode Basic,– alignSoftClipAtReferenceEnds No). Counting was carried out using STAR. Further analysis was performed for genes with a minimum of 5 reads in at least one sample. The normalization of the counts and differential expression analysis were performed using DESeq2^67^ v1.36.0 with the parameters: betaPrior=True, cooksCutoff=FALSE, independentFiltering=FALSE. Raw P values were adjusted for multiple testing using the procedure of Benjamini and Hochberg^68^. Differential gene expression was calculated by a comparison of the data of the cells exposed to different drug concentrations with the naive (control) cells. The criteria for significance were as follows: padj ≤ 0.05, |Log2FoldChange| ≥ 1, BaseMean ≥ 5. Genes taken forward for pathway enrichment were the intersection of significant DEGs between 10K drug–treated and control (naive) cells across all three HeLa clones.

### Single-cell RNA-seq

Droplet-based scRNA-seq ScRNA-seq libraries were generated using the 10X Genomics Chromium Single Cell 3’ Kit v3.1 (co-culture control experiment) and the 10x Chromium Controller (10x Genomics) according to the 10X Single Cell 3’ v2 protocol.

Final scRNA-Seq library were diluted to 4 nM, denatured, and further diluted to a final concentration of 2.8 pM for sequencing with NovaSeq SP (100 cycles) at the Weizmann Institute.

Demultiplexing and processing of the single-cell 5′ gene expression was done using the Cell Ranger ‘multi’ analysis pipeline (10x Genomics v.7.0). Cells were filtered to remove doublets (cells with two antibody tags) and negatives (no antibody tag) as well as cells with a mitochondrial gene count greater than 10%, then the singlets were assigned to the relevant samples; HeLa, Colo320DM and HT-29. For further analysis Seurat cells were further filtered and cells with less than 100 UMI counts (function subset) as well as cells in the top or bottom 1% of genes or UMI counts (function which) were removed from the analysis. Then normalization was done, dimensionality reduction was performed using PCA (RunPCA). The functions RunUMAP and FindNeighbors were applied using 10 principal components for HeLa clone 1, 20 for Colo320DM. The clustering was performed with a resolution of 0.6 (FindClusters) for HeLa and 1 for Colo320DM. Then for all three cell-lines Function VariableFeaturePlot was run with top 20 most variable genes printed for Colo320DM and HeLa clone 1. PCA and variable feature plot were run on R version 4.4.1.

### Tandem-Mass spectrometry

#### Sample preparation

HeLa clone 1 and 2 cell pellets were lysed with 5% SDS in 50 mM Tris-HCl. Lysates were incubated at 96 °C for 5 min, followed by six cycles of 30 s of sonication (Bioruptor Pico, Diagenode, USA). Protein concentration was measured using the BCA assay (Thermo Scientific, USA) and a total of 50 μg protein was reduced with 5 mM dithiothreitol and alkylated with 10 mM iodoacetamide in the dark. Phosphoric acid was added to the lysates to a final concentration of 1.2% and 90:10% methanol/50 mM ammonium bicarbonat. Each sample was then loaded onto S-Trap 96-well plate(Protifi, USA). Samples were then digested with trypsin (1:50 trypsin/protein) for 1.5 h at 47 °C. The digested peptides were eluted using 50 mM ammonium bicarbonate; trypsin was added to this fraction and incubated overnight at 37 °C. Two more elutions were made using 0.2% formic acid and 0.2% formic acid in 50% acetonitrile. The three elutions were pooled together and vacuum-centrifuged to dry. Samples were kept at −80 °C until analysis.

#### Liquid chromatography

ULC/MS grade solvents were used for all chromatographic steps. Each sample was loaded using split-less nano-Ultra Performance Liquid Chromatography (10 kpsi M-Class Acquity; Waters, Milford, MA, USA). The mobile phase was: A) H2O + 0.1% formic acid and B) acetonitrile + 0.1% formic acid. Desalting of the samples was performed online using a reversed-phase Symmetry C18 trapping column (180 µm internal diameter, 20 mm length, 5 µm particle size; Waters). The peptides were then separated using a T3 HSS nano-column (75 µm internal diameter, 250 mm length, 1.8 µm particle size; Waters) at 0.35 µL/min. Peptides were eluted from the column into the mass spectrometer using the following gradient: 4% to 30%B in 105 min, 30% to 90%B in 10 min, maintained at 90% for 7 min and then back to initial conditions.

#### Mass Spectrometry

The nanoUPLC was coupled online through a nanoESI emitter (10 μm tip; New Objective; Woburn, MA, USA) to a quadrupole orbitrap mass spectrometer (Exploris 480, Thermo Scientific) using a FlexIon nanospray apparatus (Proxeon). Data was acquired in data dependent acquisition (DDA) mode, using a 2 sec cycle. MS1 resolution was set to 120,000 (at 200m/z), mass range of 380-1500m/z, AGC of 200% and maximum injection time was set to 50 msec. MS2 resolution was set to 15,000, quadrupole isolation 1.4m/z, AGC of 75%, dynamic exclusion of 40 sec and auto maximum injection time.

#### Data processing

Raw data was processed with MetaMorpheus v1.0.2. The data was searched against the human Uniprot proteome database (xml version including known PTMs, downloaded in January 2023) appended with common lab protein contaminants and the following modifications: Carbamidomethylation of C as a fixed modification and oxidation of M as a variable one. Quantification was performed using the embedded FlashLFQ and protein inference algorithms. The LFQ (Label-Free Quantification) intensities were calculated and used for further calculations using Perseus v1.6.2.3. Decoy hits were filtered out. The LFQ intensities were log2 transformed and only proteins that had at least 2 valid values in at least one experimental group were kept. The remaining missing values were imputed. A student’s t-test was performed to identify the proteins that are differentially expressed. The mass spectrometry proteomics data have been deposited to the ProteomeXchange Consortium via the PRIDE^69^ partner repository with the dataset identifier PXD068337.

## Data Availability

Bulk RNA, single-cell RNA, nanopore, and short read paired-end DNA sequences, and mass spectrometry data will be made available upon request.

## Acknowledgments

RNA-seq analysis was performed with advice from Inbal Nachman at the Crown Genomics Institute of the Nancy and Stephen Grand Israel National Center for Personalized Medicine, Weizmann Institute of Science. Tal Bigdary from the Weizmann Institute of Science graphics unit helped with graphic design. O.S. is supported by the following research grants: the Center for New Scientists at the Weizmann Institute of Science, the ISRAEL SCIENCE FOUNDATION (grant No. 686/22), the Research Career Development Award from the Israel Cancer Research Foundation (grant No. 24-205-RCDA), the Moross Integrated Cancer Center, the Dr. Barry Sherman Institute for Medicinal Chemistry, the Minerva Stiftung - Research Grant (grant number 146552), the Flight Attendant Medical Research Institute, Florida, the Kekst Family Institute for Medical Genetics at The Weizmann Institute of Science, the Shimon and Golde Picker Annual Grant, the Weizmann - EKARD Institute for Cancer Diagnosis Research, and the Weizmann - Crown Human Genome Center.

## Contributions

S.M., I.L.D., V.N.K. and O.S. conceived the project. S.M., I.L.D. and V.N.K. performed and analysed all experiments with help from G.K., K.H, O.B., and M.C.. M.G. performed DNA sequencing analysis, which was supervised by A.G.H.. R.N. and I.A. supported imaging computational analysis. Y.R. and G.S. helped with bulk RNA-seq analysis. M.K. and G.S. helped with mass-spectrometry. R.R. helped with simulations and statistics. D.L. supported single-cell RNA-Seq analysis. N.O. performed TISSUE-FISH experiments from neuroblastoma patients and E.B. procured the samples. S.P., N.K. and Z.S. performed the short-read WGS. S.M. and O.S. wrote the manuscript with input from all co-authors.

## Ethics declarations

A.G.H. is a founder and stock holder in Econic Biosciences. The other authors declare no competing interests.

